# Multi-omics analyses of *MEN1* missense mutations identify disruption of menin-MLL and menin-JunD interactions as critical requirements for molecular pathogenicity

**DOI:** 10.1101/2022.03.22.485279

**Authors:** Koen M.A. Dreijerink, Ezgi Ozyerli-Goknar, Stefanie Koidl, Ewoud J. van der Lelij, Priscilla van den Heuvel, Jeffrey J. Kooijman, Martin L. Biniossek, Kees W. Rodenburg, Sheikh Nizamuddin, H.T. Marc Timmers

## Abstract

Loss-of-function mutations of the multiple endocrine neoplasia type 1 (*MEN1*) gene are causal to the MEN1 tumor syndrome, but they are also commonly found in sporadic pancreatic neuroendocrine tumors and other types of cancers. The *MEN1* gene product, menin, is involved in transcriptional and chromatin regulation, most prominently as an integral component of KMT2A/MLL1 and KMT2B/MLL2 containing COMPASS-like histone H3K4 methyltransferase complexes. In a mutually exclusive fashion, menin also interacts with the JunD subunit of the AP-1 and ATF/CREB transcription factors. After *in silico* screening of 253 disease-related *MEN1* missense mutations, we selected a set of nine menin mutations in surface-exposed residues. The protein interactomes of these mutants were assessed by quantitative mass spectrometry, which indicated that seven of the nine mutants disrupt interactions with both MLL1/2 and JunD complexes. Interestingly, we identified three missense mutations, R52G, E255K and E359K, which predominantly reduce the interaction with MLL1 compared to JunD. This observation was supported by a pronounced loss of binding of the R52G, E255K and E359K mutant proteins at unique MLL1 genomic binding sites with less effect on unique JunD sites. These findings support the general importance of the menin-MLL1 and menin-JunD interactions in *MEN1* gene-associated pathogenic conditions.

## Introduction

Mutations of the multiple endocrine neoplasia type 1 (*MEN1*) tumor suppressor gene are found in patients with the MEN1 syndrome (Brandi *et al*, 2020; Chandrasekharappa *et al*, 1997). MEN1 is characterized by the occurrence of parathyroid, pituitary and pancreatic neuroendocrine tumors (PanNETs). Sporadic PanNETs frequently harbor inactivating somatic *MEN1* mutations (Jiao *et al*, 2011). In addition, in recent years exome and whole genome sequencing studies have revealed *MEN1* gene mutations in many types of cancer, such as adrenocortical, uterine, breast and other cancers (Hoadley *et al*, 2018). The *MEN1* gene acts as a classic tumor suppressor gene in endocrine tissues: loss of function results in tumorigenesis. In other tissues, such as the hematopoietic system, the *MEN1* gene has pro-oncogenic activity and protein-protein interactions of the *MEN1* product, menin, are emerging as therapeutic targets (see below).

Menin is involved in transcriptional regulation as an intermediary protein linking transcription factors to co-activator and co-repressing proteins (Dreijerink *et al*, 2017b). Most notably, menin has been reported as an integral component of mixed-lineage leukemia MLL1 and MLL2 (official names: KMT2A and KMT2B) containing COMPASS-like histone H3K4 methyltransferase complexes (Hughes *et al*, 2004). Loss of *Men1*-dependent histone H3K4 trimethylation contributes to *Men1*-related pNET development in mice (Lin *et al*, 2011). Menin was co-crystallized with MLL1, the AP-1 transcription factor member JunD or a MLL1-PSIP1 (PC4 and SFRS1 interacting protein 1, also known as LEDGF, lens epithelium-derived growth factor) heterodimer (Huang *et al*, 2012). The interaction of menin with full length JunD was first reported in yeast two-hybrid and *in vitro* pull-down experiments (Agarwal *et al*, 1999). Menin contains a deep pocket that can bind short peptides of MLL1 or of JunD in a similar and mutually-exclusive manner (Huang *et al*., 2012). We performed affinity purification of menin-containing protein complexes followed by quantitative mass spectrometry, which confirmed that menin is present both in MLL1 and MLL2 containing COMPASS-like complexes and in complexes of JunD and related proteins (van Nuland *et al*, 2013).

The observation that menin is a critical cofactor for the subset of leukemias driven by chromosomal translocations involving *MLL1/KMT2A* motivated the development of menin-MLL inhibitors, which all target the MLL1-binding pocket of menin (Grembecka *et al*, 2012; Ozyerli-Goknar *et al*, 2021; Yokoyama *et al*, 2005). Several menin-MLL inhibitors display potent anti-leukemic activities in preclinical mouse models for MLL-rearranged and NPM1 mutant acute leukemias (Dzama *et al*, 2020; Klossowski *et al*, 2020; Krivtsov *et al*, 2019; Uckelmann *et al*, 2020). At present, several clinical trials investigate menin-MLL inhibitors in relapsed acute leukemias with promising early results (Issa *et al*, 2021).

The paradoxical roles of menin as a tumor suppressor for endocrine tissues and a pro-oncogenic factor for MLLr-and NPM1c-driven acute leukemias encourage a better understanding of menin function. Whereas menin inhibition provides a therapeutic perspective for certain hemopoietic malignancies, a rescue of menin function could be of therapeutic value for patients with endocrine tumors. At present, there are many unknowns with regard to loss of *MEN1* function and the subsequent events leading to endocrine tumorigenesis. Most *MEN1* gene mutations are considered to disrupt menin function entirely: nonsense *MEN1* gene mutations are predicted to result in degradation of the *MEN1* mRNA or to result in insoluble menin proteins (Brandi *et al*., 2020; Zetoune *et al*, 2008). Moreover, *MEN1* missense mutations have been reported to yield unstable protein products that are targeted for proteasomal degradation (Canaff *et al*, 2012; Yaguchi *et al*, 2004).

In this study, we sought to identify disease-related functionally impaired *MEN1* mutated forms of menin using *in silico* methods based on crystal structures of free menin and of menin-containing complexes. Such mutant forms of menin could clarify the critical pathogenic requirements for non-MEN1 conditions in terms of protein-protein interactions for loss of menin function. By using ectopic expression, we carried out quantitative proteomic studies of a selection of mutant menin proteins in HeLa cells. We identify a set of disease-associated menin mutants with differential effects on MLL1 and JunD interactions, which were validated by analyzing genome localizations of wild type (WT) and the mutant forms of menin, as well as JunD, MLL1 and histone modifications.

## Materials and methods

### Identification of disease-associated MEN1 mutations

A list of *MEN1* gene missense mutations was compiled using data from the Human Gene Mutation Database (HGMD http://www.hgmd.cf.ac.uk/ac/index.php) for *MEN1* mutations and the Catalogue of Somatic Mutations in Cancer (COSMIC, http://cancer.sanger.ac.uk/cosmic) for somatic *MEN1* mutations, complemented with data from the literature (Agarwal *et al*., 1999; Canaff *et al*., 2012; Huang *et al*., 2012)(Supplementary Table 1).

### In silico analysis of MEN1 missense mutation-derived menin proteins

The PDB file of menin (3U84, reduced to monomer, containing 550 amino acids of the 610 amino acid menin protein) was analyzed using Swiss-PdbViewer 4.1.0. Menin protein products resulting from *MEN1* mutations were assessed based on the following criteria: surface accessibility, predicted charge and polarity changes due to the mutations, presence of hydrogen bonds as well as salt bridges at the site of the mutations (using VMD 1.9.2 (http://www.ks.uiuc.edu/Research/vmd/)). For every mutation, the number of rotamers, which showed clashes within the tertiary protein structure was counted (Supplementary Table 1). When using default software settings, the absence of clashing rotamers (using default software settings) due to the *MEN1* mutations was defined as stable, less than 100% clashing rotamers as potentially stable and 100% clashing rotamers due to the mutation as predicted to result in unstable menin protein.

The PDB files of the menin-MLL1 complex (3U85), menin-MLL1-LEDGF complex (3U88) and menin-JunD complex (3U86) were acquired from RCSB (www.rcsb.org) for analysis of the effects of *MEN1* mutations on the reported interactions of menin with MLL1, PSIP1/LEDGF or JunD. Mutations in the reported MLL1, PSIP1/LEDGF and JunD interaction surfaces of menin were described by selecting amino acids in a range of 5 Å from a known interaction site and verifying if any known mutations were present within this 5 Å-radius. Only mutations on the surface and affecting an amino acid capable of forming H-Bonds were considered for their relevance for the predicted interaction surface.

### Projection of menin mutations

Both structures of the menin-JunD (ID: 3U86) and menin-MLL1-LEDGF (ID: 3U88) complexes were superimposed by sequence alignment with matchmaker and amino acids were highlighted using Chimera (version 1.13.1) (Pettersen *et al*, 2004).

### Plasmids and cell lines

The *MEN1* cDNA was cloned into the Gateway system using the pENTR directional TOPO cloning kit to enable C-terminal tagging (Life technologies, Thermo Fisher Scientific, MA, USA). *MEN1* gene mutations were introduced using site-directed mutagenesis essentially as described by the Quikchange strategy (Agilent, CA, USA) into the MEN1 cDNA in a pCDNA3.1 vector for transient expression, as well as in a pENTR-menin plasmid for C-terminal tagging with GFP via GATEWAY recombination with the pCDNA5_FRT_TO_C-GFP plasmid (van Nuland *et al*., 2013). All plasmids were validated by DNA sequence analysis and primer sequences are available on request.

Human embryonic kidney (HEK) 293T and human cervical carcinoma HeLa cells were cultured in standard growth medium (DMEM, 10% FBS, 100 U/ml penicillin, 150 ug/ml streptomycin and 2 mM glutamine).

### Expression of menin (mutant) proteins

HEK 293T cells were transiently transfected with 600 ng DNA in a 12-well format using the PEI transfection reagent. After 48 hours, cells were lysed in Laemmli sample buffer. Total cell lysates were analyzed through immunoblotting using anti-menin (A300-105A, Bethyl) and anti-alpha tubulin (CP06 Calbiochem) antibodies respectively.

For doxycycline-inducible expression in stable cell lines MEN1-GFP expressing cDNAs were chromosomally integrated in HeLa cells using the Flip-in system, essentially as described (van Nuland *et al*., 2013). After 24 hours of induction using 1 μg/ml doxycycline, GFP expression was verified using immunoblotting using GFP (JL-8-Clontech) and α-tubulin (CP06 Calbiochem) antibodies. Nuclear and cytoplasmic extracts were prepared for GFP-affinity purification coupled to mass spectrometry analyses as described before (van Nuland *et al*., 2013).

### Quantitative mass spectrometry of the menin interactome

GFP-affinity purification, sample preparation and data analysis were performed as reported (Nizamuddin *et al*, 2021; Spruijt *et al*, 2013; van Nuland *et al*., 2013). Briefly, HeLa cell harboring menin-GFP cDNAs were grown in 15 15-cm dishes until subconfluency (approximately 300 million cells in total) and induced with 1 ug/mL doxycycline for 24h. Cells were harvested by dislodging with trypsin and cell pellets washed with cold PBS (Gibco, #10010-015). Cell pellet was re-suspended in 5 packed-cell volumes (PCVs) of ice-cold Buffer A (10 mM Hepes-KOH pH 7.9, 1.5 mM MgCl_2_, 10 mM KCl), incubated for 10 min on ice and then centrifuged at 400 *g* and 4°C for 5 min. Supernatants was aspirated and cells were lysed in 2 PCVs Buffer A containing 1x CPI (Roche, #11836145001), 0.5 mM DTT and 0.15 % NP40. The suspension was homogenized in Dounce homogenizer followed by centrifugation at 3,200 *g* and 4°C for 15 min. Supernatant and pellet contain cytoplasmic and nuclear fractions, respectively. The nuclear pellet was washed gently with 10 volumes of Buffer A containing 1x CPI (Roche, #11836145001), 0.5 mM DTT and 0.15 % NP40 and centrifuged for 5 min at 3,200 *g* at 4°C min. Nuclear proteins were extracted by 2 PCVs volumes of high salt Buffer B (420 mM NaCl, 20 mM Hepes-KOH pH 7.9, 20% v/v glycerol, 2 mM MgCl2, 0.2 mM EDTA, 0.1 % NP-40, 1x CPI, 0.5 mM DTT) during gentle agitation at 4°C for 1.5 h. Both the nuclear and cytoplasmic extracts were centrifuged at 3,200 *g* and 4°C for 60 min. Supernatants were collected and protein concentration was measured by Bradford assay.

1 mg of nuclear or 3 mg of cytoplasmic extract was used for GFP-affinity purification as described(Spruijt *et al*., 2013). In short, protein lysates were incubated in binding buffer (20 mM Hepes-KOH pH 7.9, 300 mM NaCl, 20% glycerol, 2 mM MgCl2, 0.2 mM EDTA, 0.1% NP40, 0.5 mM DTT and 1x Roche protease inhibitor cocktail) on a rotating wheel at 4° C for 1 h in triplicates with GFP-Trap agarose beads (#gta-200, Chromotek) or control agarose beads (Chromotek). The beads were washed two times with binding buffer containing 0.5% NP-40, two times with PBS containing 0.5% NP-40, and two times with PBS. On-bead digestion of bound proteins was performed overnight in elution buffer (100 mM Tris-HCl pH 7.5, 2 M urea, 10 mM DTT) with 0.1 μg/ml of trypsin at RT and eluted tryptic peptides were bound to C18 stage tips (ThermoFischer, USA) prior to mass spectrometry analysis.

Tryptic peptides were eluted from the C18 stage tips in H_2_O:acetonitrile (35:65) and dried. Samples were analyzed by nanoflow-LC-MS/MS with a Q-ExactivePlus mass spectrometer (Thermo Fischer Scientific) coupled to an Easy nano-LC 1000 HPLC (Thermo Fisher Scientific) in the tandem mass spectrometry mode with a 90 min total analysis time. The flow rate was 300 nl/min, buffer A was 0.1 % (v/v) formic acid and buffer B was 0.1 % formic acid in 80 % acetonitrile. A gradient of increasing organic proportion was used in combination with a reversed phase C18 separating column (2 μm particle size, 100 Ǻ pore size, 15 cm length, 50 μm i.d., Thermo Fisher Scientific). Each MS scan was followed by a maximum of 10 MS/MS scans in the data-dependent mode. Blank samples were run between each set of three samples to minimize carry over.

The raw data files were analyzed with MaxQuant software (version 1.5.3.30) using Uniprot human FASTA database (Spruijt *et al*., 2013; Tyanova *et al*, 2016). Label-free quantification values (LFQ) and match between run options were selected. Intensity based absolute quantification (iBAQ) algorithm was also activated for subsequent relative protein abundance estimation (Schwanhäusser *et al*, 2011). The obtained protein files were analyzed by Perseus software (MQ package, version 1.6.12), in which contaminants and reverse hits were filtered out (Tyanova *et al*., 2016). Protein identification based on non-unique peptides as well as proteins identified by only one peptide in the different triplicates were excluded to increase protein prediction accuracy.

For identification of the bait interactors LFQ intensity-based values were transformed on the logarithmic scale (log2) to generate Gaussian distribution of the data. This allows for imputation of missing values based on the normal distribution of the overall data (in Perseus, width = 0.3; shift = 1.8). The normalized LFQ intensities were compared between grouped GFP triplicates and non-GFP triplicates using 1% as the permutation-based false discovery rate (FDR) in a two-tailed t-test. The threshold for significance (S0), based on the FDR and the ratio between GFP and non-GFP samples was kept at the constant value of 2. Relative abundance plots were obtained by comparison of the iBAQ values of GFP interactors. The values of the non-GFP iBAQ values were subtracted from the corresponding proteins in the GFP pull-down and were next normalized on the menin-GFP bait protein for scaling and data representation purposes. All mass spectrometry data have been deposited to the ProteomeXchange Consortium via the PRIDE partner repository under the dataset identifier PXD031928.

### Genome localization experiments by (green)CUT&RUN

Genome localization analysis of GFP-tagged menin and GFP-tagged JunD was performed by greenCUT&RUN with the combination of enhancer-MNase and LaG16-MNase as described (Koidl & Timmers, 2021). To localize MLL1 or RNA polymerase II standard CUT&RUN (Skene *et al*, 2018) was employed using MLL1/KMT2A antibodies (Epicypher #SKU 13-2004) or 8WG16 antibodies directed against the CTD of RPB1 at 1 μg/ml. For standard CUT&RUN and greenCUT&RUN mononucleosomal *Drosophila* DNA was used as spike-in DNA for normalization purposes and sequencing libraries were prepared as described (Nizamuddin *et al*., 2021). In brief, purified DNA fragments were subjected to library preparation with NEB Next Ultra II and NEB Multiplex Oligo Set I/II as per manufacturer (New England Biolabs) protocol without size selection. For each library, DNA concentration was determined using a Qubit instrument (Invitrogen, USA) and size distribution was analyzed with Agilent Bioanalyzer chips (DNA high sensitivity assay). The 75-nucleotide paired-end sequencing reads were generated (Illumina, HiSeq 3000) with 6-32 M reads per sample (Supplementary Table 2). These NGS data have been deposited to Sequence Read Archive (https://www.ncbi.nlm.nih.gov/sra) under the accession number PRJNA772915.

### Bioinformatic analyses of genomics data

The HeLa cell datasets for H3K4me3 (ID: ENCFF063XTI), H3K4me1 (ENCFF617YCQ) and H3K27ac (ENCFF113QJM) were downloaded from ENCODE (www.encodeproject.org). The ATACseq dataset for HeLa cells was downloaded from SRA-NCBI (accession ID: SRR8171284).

The datasets generated in-house were initially passed through quality control using Trim-galore (version 0.6.3). Further reads were aligned on the human (version hg38) and *Drosophila* reference genome (BDFP5) using bowtie2 (version 2.3.4.1) with option: –dovetail –local –very-sensitive-local –no-unal –no-mixed –no-discordant -I 10 -X 700 (Langmead & Salzberg, 2012; Nizamuddin *et al*., 2021). Equal number of reads was randomly selected for WT menin and menin mutants using sambamba (version 0.6.9) and utilized in the further analysis. The correlation plot was generated using the deeptools (version 3.3.2) (Ramirez *et al*, 2014; Tarasov *et al*, 2015).

Peaks were called using HOMER with default parameters except filtering based on clonal signals was disabled using option: -C 0. Both narrow and broad peaks were called with HOMER and merged together using bedtools (version 2.27.1) except for JunD for which only narrow peaks were called (Quinlan & Hall, 2010). Reads of the control samples were normalized with SpikeIn and used to generate TagDirectories by HOMER using option: - totalReads before peak calling. To calculate normalized total reads, ratio of the SpikeIn per human reads of control and experiment were calculated and multiplied with total number of control reads. In case of H3K4me3 (ENCFF862LUQ) the coordinate of the peaks were downloaded from ENCODE. The peaks of menin (wild type) overlapping with JunD, MLL1 and H3K4me3 were identified using bedtools and classified into eight categories using in-house script which can be downloaded from https://github.com/snizam001/MEN1 (ann.R).

HOMER was used to find AP1/ATF motifs within the peaks with default parameters. In differential peak analysis, menin peaks with coverage of fold changes ≤4 or p value ≥0.001 against mutants were considered as unaffected by the menin mutations.

Heatmaps were generated using deeptools (version 3.3.2) with default parameters. All next-generation sequencing datasets have been deposited to the Sequence Read Archive (SRA) portal of the NCBI with accession ID PRJNA772915. The command-lines used for the data analysis can be downloaded from https://github.com/snizam001/MEN1.

## Results

### In silico analysis

We selected *MEN1* gene mutations from a compiled list of 253 missense mutations observed in various human cancers. These missense mutations correspond to 245 unique single amino acid changes, which were subjected to *in silico* modeling for structural changes using menin crystal structure data (Supplementary Table 1). The selection criteria included predicted protein stability and predicted disruption of menin-MLL1 and/or menin-JunD interactions. Of these 245 *MEN1* missense mutations 80 were expected to affect protein stability, because the mutation did not yield any fitting rotamers in our modeling. For 47 missense mutations the decrease in protein stability was predicted from a loss of internal hydrogen bonds. 86 of the 253 studied mutations appeared to be potentially pathogenic for unknown reasons other than losing internal bonds or directly altering protein-protein interactions with the MLL1 and JunD complexes. Because of their proximity to the mapped interaction surfaces of menin and MLL1 and JunD respectively, six missense mutations were predicted to potentially directly alter the menin binding to MLL1 or JunD (Figure 1A and 1B).

**Figure 1.**
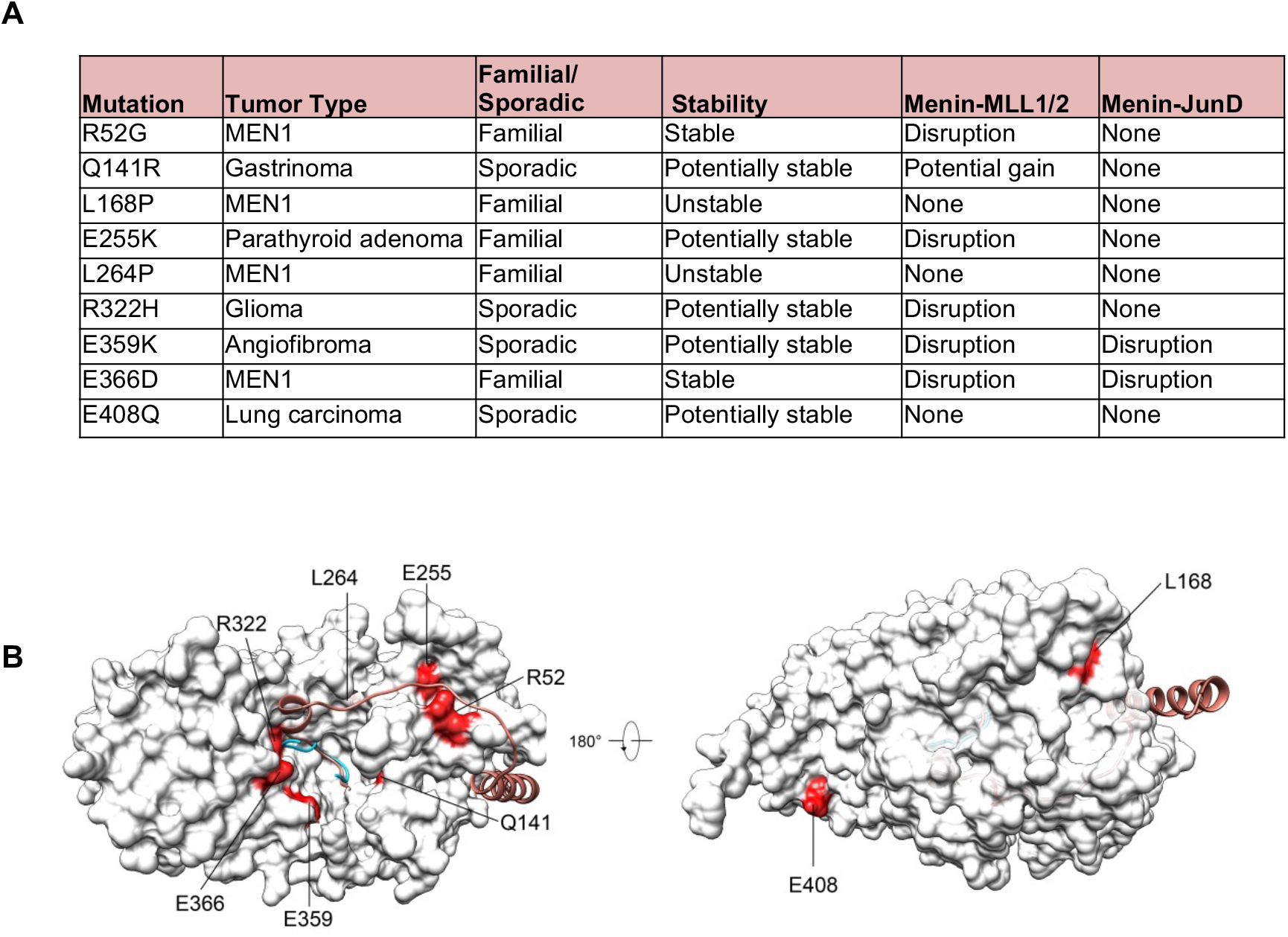
Selection of *MEN1* missense mutation-derived forms of menin. Panel A. Nine disease-relevant *MEN1* missense mutations were selected based on their predicted stability *in silico* and ability to bind to MLL1 and/or JunD. The predicted effects of the mutations are listed. Panel B. The nine *MEN1* mutations are projected on the menin structure.

For the R52G and E255K mutants, disruption of the interaction of menin with MLL1 when in complex with LEDGF was predicted. The R322H, E359K and E366D mutations could disrupt both the interaction of menin with MLL1 and with the MLL1-LEDGF complex, while E359K and E366D were predicted to disrupt the interaction of menin with JunD in *silico*. The Q141R mutation was selected as it was considered to be a stable protein and to potentially result in a gain of interaction of menin with MLL1, when in complex with LEDGF.

### Characteristics and transient ectopic expression of selected MEN1 mutations

The selected *MEN1* mutants are associated with different human endocrine or cancer pathologies (Figure 1A). The R52G mutation was first identified in a MEN1 patient who developed primary hyperparathyroidism and a PanNET (Hou *et al*, 2011). Menin mutations L168P, L264P and E366D have been identified in MEN1 families (Bartsch *et al*, 1998; Jäger *et al*, 2006; Poncin *et al*, 1999). The Q141R mutation was identified in a sporadic gastrinoma without loss of heterozygosity LOH at the wild type (WT) locus, which is common in gastrinomas (Wang *et al*, 1998). Menin E255K has been found in a kindred with familial isolated hyperparathyroidism, a condition considered to be a variant phenotype of MEN1 (Teh *et al*, 1998). The R322H mutation was found in a sporadic glioma (TCGA-CS-5394-01). The mutation resulting in menin E359K was found in a sporadic angiofibroma, a skin tumor type known to be associated with MEN1, without LOH at the WT locus (Böni *et al*, 1998; Darling *et al*, 1997). The *MEN1* gene mutation resulting in menin E408Q was derived from a non-small cell lung cancer sample (TCGA-22-4607-01).

Based on this and on our *in silico* analysis, we selected missense menin mutants R52G, E255K, R322H, E359K, E366D and Q141R for further functional studies. In addition, we included the L168P and L264P mutations, which are predicted to be unstable, as well as the E408Q mutation from a non-endocrine tumor as a control. E408Q was predicted to be stable and have no effect the menin-MLL1 and menin-JunD interactions as it is located far from the binding pocket (Figure 1B).

Next, we determined the stability of the selected mutant forms of menin by ectopic expression as described before (Yaguchi *et al*., 2004). The nine different *MEN1* mutants, along with a vector control, were introduced into a CMV-driven expression plasmids, which were used for transient transfection of HEK 293T cells in order to assess their expression levels in total cell lysates as measured by immunoblotting using menin antibodies. Please note that the signal in the pCDNA control results from endogenous menin. As shown in Figure 2A, immunoblotting of transfected cell lysates confirmed stability of the selected mutations. As expected, the L168P and L264P menin proteins were expressed at lower levels compared with WT menin, indicating a reduced protein stability as noted before (Dreijerink *et al*, 2006).

**Figure 2.**
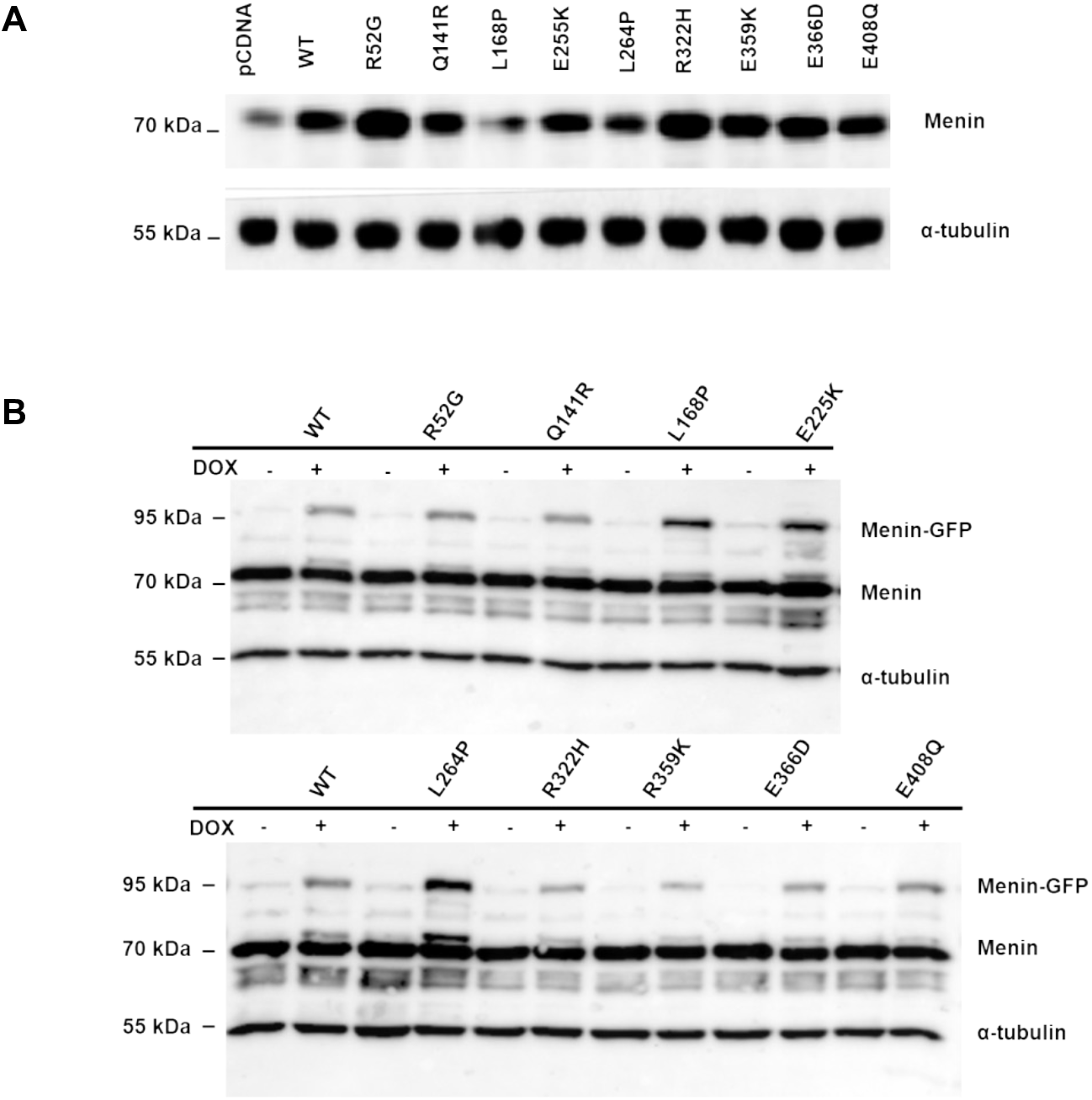
Expression levels of menin proteins after transient transfections of HEK293 cells and or stable integration in stable HeLa cell lines. Panel A. Protein stability was assessed by ectopic expression of *MEN1* WT and mutant cDNA in HEK293T cells and subsequent immunoblotting using menin antibodies or α-tubulin antibodies as a loading control. Panel B. Immunoblots upon doxycycline induction of stable HeLa cell lines showing expression of GFP-tagged menin using menin antibodies or α-tubulin antibodies as a loading control.

### Quantitative mass spectrometry of stably expressed menin mutants

We proceeded to assess the menin interactome using C-terminal GFP fusions of the menin mutants by quantitative mass spectrometry after GFP-based immunoaffinity purification. For this purpose, *MEN1* WT and mutant cDNA fusions were expressed in stably transfected HeLa cell lines in an inducible manner. Immunoblot analyses of total lysates with menin antibodies indicated inducible and similar expression of the GFP-fused WT and mutant menin proteins (Figure 2B) with the exception of L168P and L264P. In contrast to the HEK293T transient transfection experiment (Figure 2A), GFP-fusions of these two mutants are expressed to higher levels than WT menin. We noted that the menin-GFP proteins are expressed to lower levels than endogenous menin (Figure 2B). Fluorescence microscopy confirmed the nuclear presence of the GFP-menin WT protein as well as the entire set of mutants with the exception of menin L168P and L264P, which are predominantly present in the cytoplasm (data not shown).

Quantitative mass spectrometry of purified WT menin-GFP identified all MLL1/MLL2 complex members (DPY30, KMT2A, KMT2B, WDR5, ASH2L, RBBP5, HCFC1, HCFC2, and PSIP1) as significant interactors (indicated by red dots in Figure 3A). Also, a weak interaction with OGT was observed as reported before (van Nuland *et al*., 2013). In addition, JunD and related proteins ATF7 and FOS were found as significant interactors (marked by blue dots in Figure 3A). The AP1 and ATF complex members FOSL2, ATF2 and ATF3 did not pass our stringent significance criteria. Based of LFQ values we determined the stoichiometries of interacting proteins relative to the WT and mutant menin-GFP baits as shown in Figure 3A and 3B. As expected, GFP-purification of the unstable L168P and L264P mutants, which are mostly cytoplasmic, did not yield any significant MLL1/MLL2 complex or JunD-related interactions (Figure 3B). Menin Q141R displayed strongly reduced, but significant interactions with MLL1/MLL2 or JunD and the interactions displayed by menin E408Q were very similar to WT menin (Figure 3B). Immunopurification of menin mutants R52G, E255K and E359K resulted mostly in loss of MLL1/MLL2 subunits with a smaller effect on JunD-related interaction (Figure 3B). Interestingly, the E366D and R322H mutants, which were predicted to disrupt the menin-MLL interactions, retained MLL1/MLL2 complex interactions. In contrast to the *in silico* predictions, JunD binding was retained entirely by the E366D mutation and at least partially in menin R322H and E359K complexes (Figure 3B). Relative MLL1 and JunD binding of all menin mutants compared with WT menin is summarized in Figure 3C. The Volcano plots of the menin mutants are presented in Supplementary Figure 1. Next, we compared the cytoplasmic interactions of the set of GFP-menin proteins. Among the top 10 interactors of cytoplasmic WT menin are ubiquitin, the ubiquitin E3 ligase RNF126, the ubiquitin-chain binders RAD23A/B and VCP, and protein chaperones (HSPA8, HSPA1A/B, DNAJB1) (Figure 4A). This indicates that newly-synthesized WT menin-GFP fusion is subjected to ubiquitin-dependent proteosomal turnover in the cytoplasm, which is consistent with previous observations of transiently expressed WT and mutant menin proteins (Yaguchi *et al*., 2004). While in this earlier work CHIP(STUB1) was identified as the responsible ubiquitin-protein E3 ligase in HEK293T cells, our results suggest that RNF126 is responsible for menin turnover in HeLa cells.

**Figure 3.**
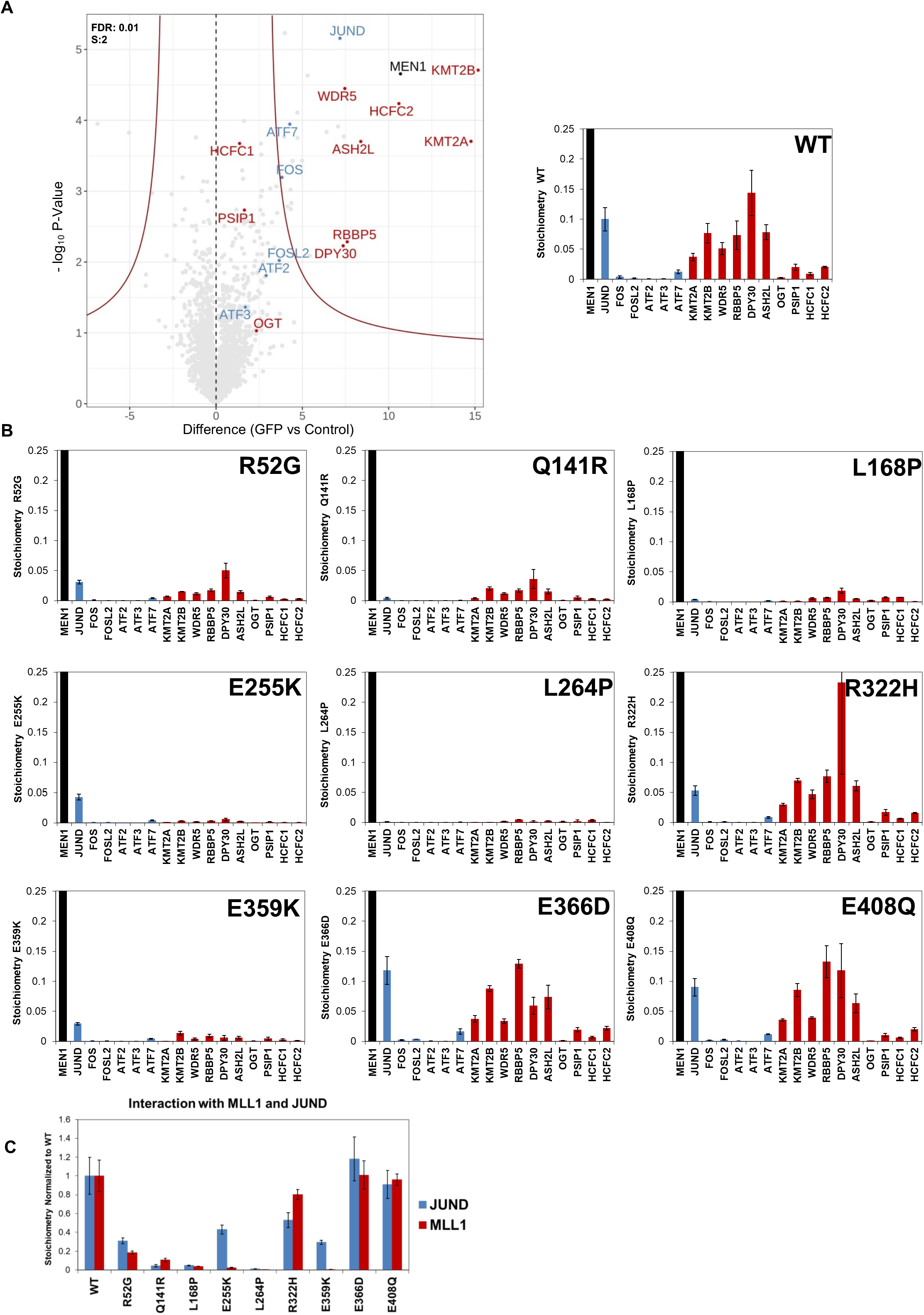
Quantitative proteomics of nuclear GFP-tagged WT menin or mutant menin expressed in HeLa cells. Panel A. Volcano and stoichiometry plots of significant interactors of WT menin-GFP isolated from nuclear extracts. Panel B. Stoichiometry plots of all mutant menin-GFP proteins for subunits of MLL1/MLL2 (red bars)- and JunD (blue bars)-containing complexes. All interactors were normalized to the menin-GFP bait. Panel C. Summary of MLL1/MLL2 complex (red bars) and of JunD complex (blue bars) interaction stoichiometries of menin mutants relative to WT menin-GFP. Results shown represent iBAQ (intensity Based Absolute Quantification) values with whiskers indicating standard deviations.

**Figure 4.**
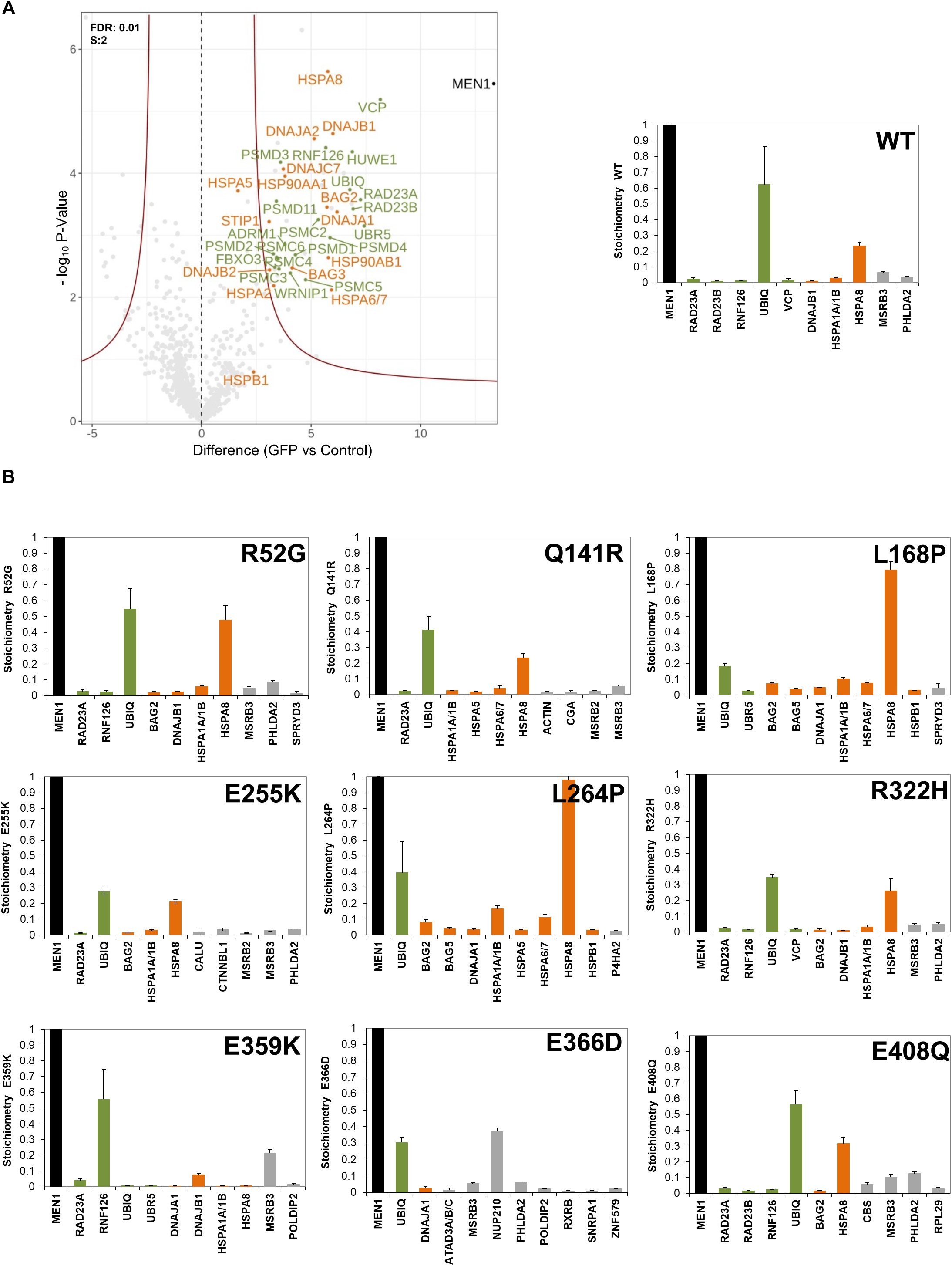
Quantitative proteomics of cytoplasmic GFP-tagged WT menin or mutant menin expressed in HeLa cells. Panel A. Volcano and stoichiometry plots for the significant interactors of WT menin-GFP from the cytoplasmic fraction. Panel B. iBAQ-based stoichiometries of the top 10 cytoplasmic interactors normalized to the WT or mutant menin-GFP bait protein. Results shown represent Intensity Based Absolute Quantification with whiskers indicating standard deviations. Please note that the MaxQuant search identified RPS27A and UBA52 as cytoplasmic menin inhibitors, but manual inspection of peptides assigned to RPS27A and UBA52 uniquely map to the ubiquitin part of these ubiquitin fusion proteins and no RPS27A nor UBA52 specific peptides could be identified.

Subsequent analyses of the cytoplasmic interactome of the stable mutations showed similar interaction patterns (Figure 4B and Supplementary Figure 2). However and as expected (Dreijerink *et al*., 2006), the two predicted unstable *MEN1* mutations resulting in menin L168P and L264P exhibited more pronounced HSPA8 and HSPA1A/B binding compared to the other menin mutants or to WT menin. HSPA8(HSC70) and HSPA1A/B(HSP70) are the most abundant protein chaperones in HeLa cells. These ATP-hydrolyzing chaperones act to assist in pulling nascent polypeptides out of the ribosome, proper *de novo* folding, assembly of protein complexes and controlling the degradation of mis-folded proteins (Finka *et al*, 2015). Interestingly, BAG2 and BAG5, which act as nucleotide-exchange factors for HSPA8 and HSPA1A/B, are also amongst the most abundant menin (mutant) interactors. In conclusion, the increased association of HSPA8A and HSPA1A/B is consistent with misfolding of the L168P and L284P menin mutants. However, WT menin and other mutants predicted to be stable also associate with chaperones and components of the controlling protein quality and abundance, which indicates that menin proteins are under strict surveillance for protein quality.

### Analyses of genome-wide binding of menin mutants and co-occupancy with JunD and MLL

In order to determine the functional relevance of the interactome results, we determined genome-wide DNA binding of the GFP-tagged menin protein and the R52G, E255K and E359K mutants, that displayed a predominant disruption of the menin-MLL1/MLL2 over the menin-JunD interactions. First, we determined the genome distribution of MLL1 and of RNA polymerase II (pol II) by CUT&RUN (Skene *et al*., 2018) using MLL1 or RPB1-CTD antibodies, respectively, and of GFP-tagged JunD by greenCUT&RUN using a combination of enhancer-MNase and LaG16-MNase as described (Koidl & Timmers, 2021). In both methods we included mononucleosomal *Drosophila* DNA as a spike-in for normalization purposes (Nizamuddin *et al*., 2021). The ENCODE HeLa cell ChIPseq datasets for the histone modifications H3K4me3, H3K4me1 or H3K27ac and for DNA accessibility determined by ATAC-seq were included. We found that WT menin is predominantly present at sites of active transcription as indicated by the presence of H3K4me3 and pol II as well as accessible DNA. Menin appeared to bind DNA either in a broad or a sharp pattern (Figure 5A). The broad pattern overlaps with MLL1 and H3K4me3, whereas a number of sharp menin peaks seemed devoid of MLL1. In order to examine this further, menin peaks were classified into eight categories on the basis of its overlap with the MLL1, JunD and H3K4me3 peaks. This resulted in eight different clusters (Figure 5B). In the largest cluster 1 (9009 peaks, 49.9%) menin binding overlaps with MLL1, H3K4me3 and JunD. These sites are also marked by H3K27ac, pol II binding and DNA accessibility indicating that menin binding correlates with promoter activity as noted before in ChIPseq experiments (Dreijerink *et al*, 2017a; Krivtsov *et al*., 2019; Uckelmann *et al*., 2020). The sharp menin peaks overlap with JunD and not always with MLL1 as indicated by cluster 3 and 4 (1776 peaks or 9.3% and 1170 peaks or 6.2%, respectively). A high percentage of the peaks in these two JunD^+^/MLL1^-^ clusters contain cognate AP1 or ATF consensus sequences (35.5% and 49.7% with AP1 motifs, and 12.33% and 15.04% with ATF motifs within cluster 3 and 4, respectively; Figure 6A). In contrast, the cluster 1 of broad menin peaks overlapping with JunD, MLL1 and H3K4me3 peaks contain fewer AP1 or ATF motifs. Menin peaks, coinciding with MLL1 and H3K4me3 presence but not JunD binding were found to harbor the fewest AP-1 binding sites (clusters 5 and 6). Similar analyses were carried out for the menin mutants R52G, E255K, E359K and menin E408Q was included as a control. As expected, menin E408Q clustered with WT menin, while the other mutations form a different cluster in the correlation analysis (Figure 6B). For global comparisons of menin-MLL or JunD interaction loss, we focused on WT menin-JunD peaks in the absence of MLL1/H3K4me3 peaks (cluster 4) and WT menin-MLL1/H3K4me3 peaks in the absence of JunD (cluster 5). We observed loss of menin R52G, E255K, E359K binding both to sites, that were co-occupied by WT menin and MLL/H3K4me3, and to sites, that were co-occupied by WT menin and JunD (Figure 6C). However, mutant menin binding to menin-MLL1/H3K4me3 sites was disrupted to a larger extent compare to menin-JunD. The difference was more pronounced in menin E255K, E359K than in menin R52G. Taken together, the genome localization data of the R52G, E255K, E359K menin mutants are consistent with the interactome data, and indicate that disruption of the interaction with the MLL1/MLL2- and JunD-containing complexes underwrite the molecular defects of disease-related mutations in *MEN1*.

**Figure 5.**
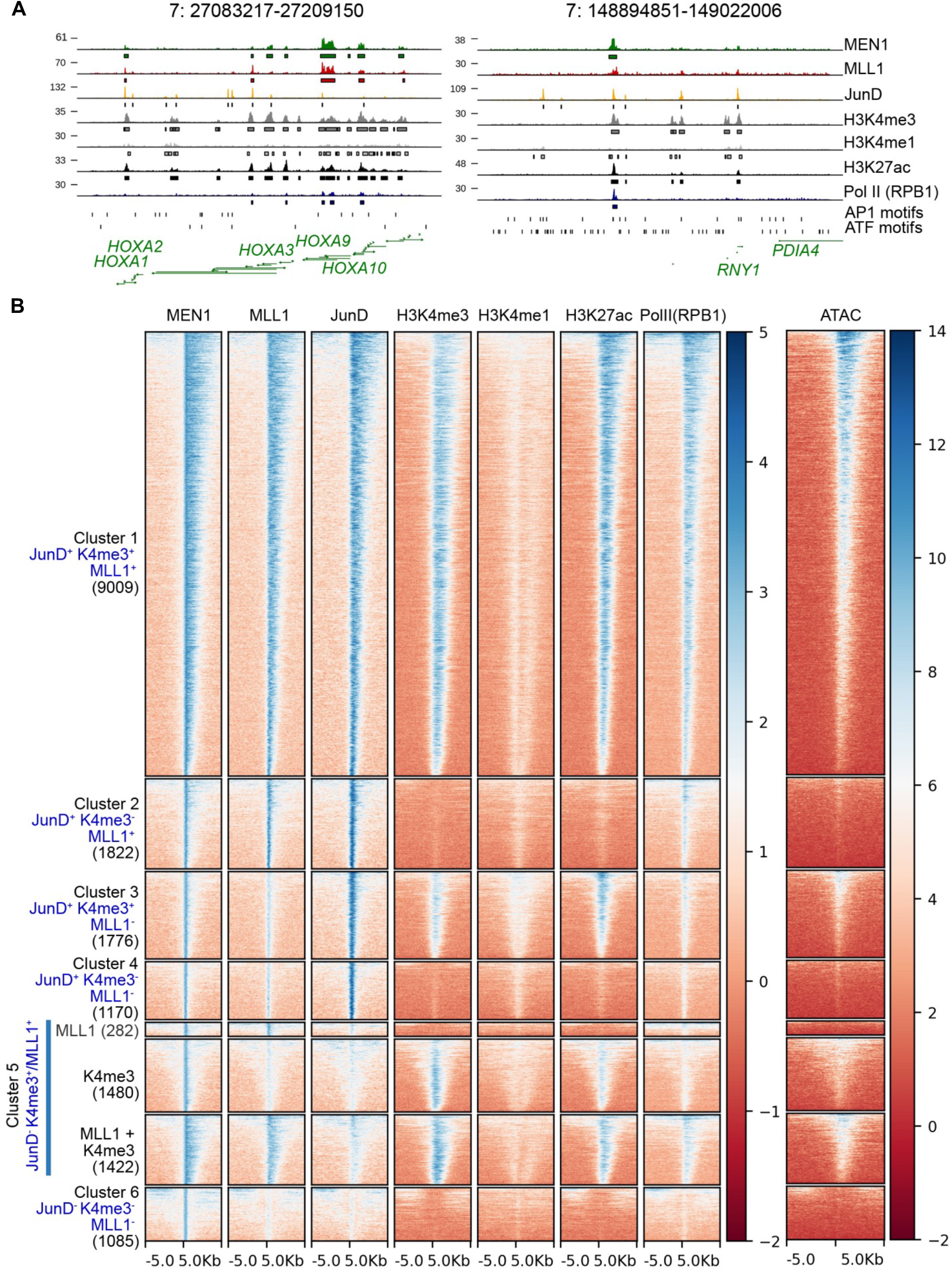
Genome-wide occupancy of WT menin, MLL1, JunD, RNA polymerase II, genome-wide presence of histone marks and DNA accessibility in HeLa cells. Panel A. Normalized coverage tracks of the menin occupied regions showing broad and narrow peaks. Panel B. Normalized heatmaps indicating menin, MLL1, JunD, H3K4me3, H3K4me1, H3K27ac and pol II peaks, as well as ATACseq results categorized according to menin, MLL1 and JunD binding. The + and - signs indicates overlap and non-overlap with the peaks of WT menin. The eight clusters are indicated on the left and the numbers between brackets indicate the total number of menin peaks in each cluster.

**Figure 6.**
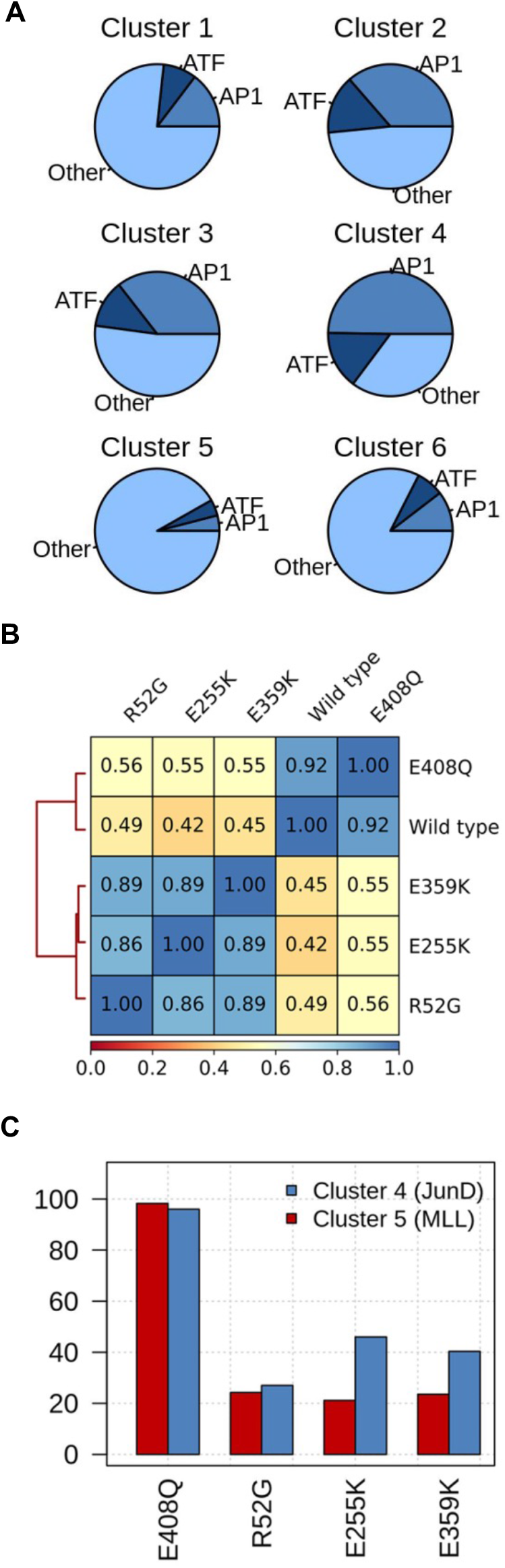
Distribution of the motifs in different clusters and genome-wide binding events of the menin mutants. Panel A. Pie charts showing the proportion of AP-1 and ATF motifs at menin-JunD sites in the different clusters from Figure 5B. Other indicates that the absence of a strong TF consensus site in this group. Panel B. Correlation of the coverage between WT menin and menin mutants. Values represent Pearson correlations of normalized coverage calculated for 10-kb bins. C. Percentages of peaks retained at menin WT bound sites by greenCUT&RUN mapping using the selected menin mutations. In blue, retained binding at menin-JunD sites without MLL1/H3K4me3 is shown. In red, retained binding of mutant forms of menin at WT menin-MLL1/H3K4me3 sites without JunD is shown.

## Discussion

Understanding the molecular pathology caused by *MEN1* gene mutations may help to elucidate the pathophysiology of MEN1-associated tumorigenesis. Our study started with an *in silico* screening approach based on crystal structures of menin, combined with immunoaffinity purification and quantitative mass spectrometry experiments to identify functionally impaired *MEN1* gene missense mutation-derived proteins. Subsequently, we functionally validated our observations by analyzing the genome localization of selected mutant forms of menin. Using these methods, we successfully identified menin missense mutations (R52G, Q141R, L168P, E255K, L264P, R322H and E359K) that directly result in loss of the menin-MLL1/2 complex and of menin-JunD interactions, indicating that disruption of this interaction is the most critical link in the molecular pathogenicity of menin protein function loss. While R52G, L168P and L264P were identified in MEN1 patients, the Q141R, E255K and E359K mutations were isolated from patients with pathologies related to MEN1 (Figure 1A).

The R52G and E359K mutants have not been analyzed in protein interaction studies before. R52G was found in a MEN1 family of three generations (Hou *et al*., 2011). The E359K mutation was found in an angiofibroma, a skin tumor type known to be affected by *MEN1* gene mutations (Böni *et al*., 1998; Darling *et al*., 1997). The E255K mutation has been studied more extensively. This mutation was reported in a familial isolated hyperparathyroidism family and found to co-segregate with the occurrence of parathyroid adenomas in this family. Loss of the WT allele was confirmed in parathyroid adenoma tissue. E255K has been described in several studies on properties of the menin protein. It was found to have a half-life similar to WT menin supporting the observation, that it is a stable *MEN1* gene missense mutation (Yaguchi *et al*., 2004). Apart from a previous report from our group in which we showed reduced interaction of menin E255K with the estrogen receptor alpha ligand binding domain in a yeast two hybrid set-up, the menin E255K mutation has not been studied for its ability to interact in MLL1/2 complexes or with JunD before (Dreijerink *et al*., 2006). Interestingly, within the three-dimensional structure of menin, residues R52 and E255 are in close proximity to each other (Figure 7). Structural analyses indicate that R52, E255 and E359 are all part of the interaction surface with MLL1, and therefore, we expected loss of MLL1 interaction of these mutants. In addition to binding MLL1, E359 is also involved in the interaction with JunD. The incomplete loss of menin E359K-JunD binding (Figure 3B) could be due to additional menin-JunD binding surfaces. For E255K and E359K we also find menin-JunD interactions are less affected compared to MLL1/MLL2 interactions. These observations are in line with the lack of clinical correlation of endocrine tumorigenesis and JunD function, for example the absence of *JUND* gene mutations in MEN1-related endocrine tumor types. This indicates that the loss of incorporation of mutant forms of menin into MLL1 and MLL2 histone methyltransferase complexes is a central feature of stable menin missense mutations in MEN1-related endocrine tumorigenesis.

**Figure 7.**
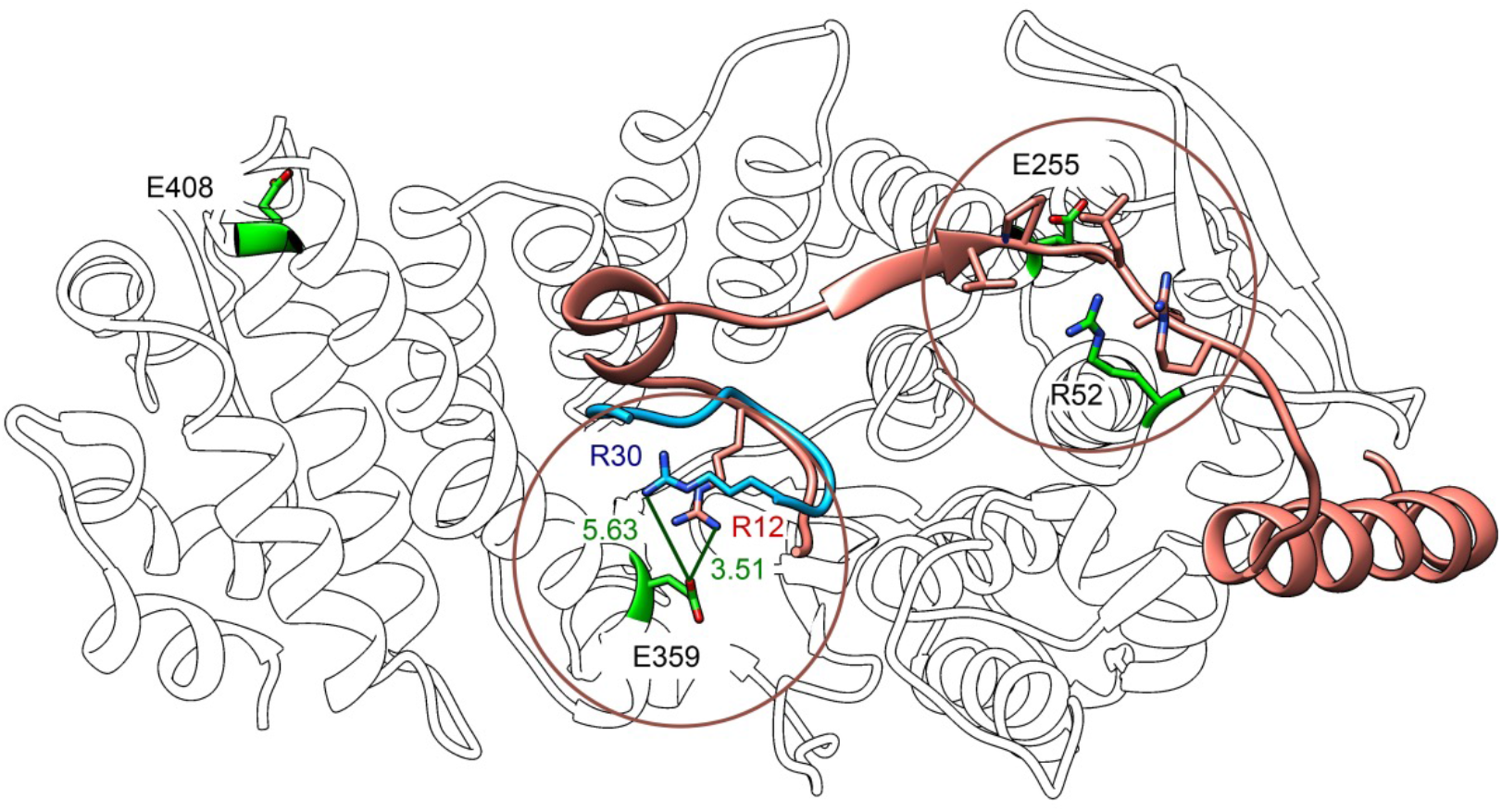
Projection of the selected *MEN1* gene mutations menin R52G, E255K, E359K and E408Q (in green) on the tertiary structure of WT menin. In brown, the position of MLL1 peptide used for the *in silico* analyses is shown. The blue peptide indicates the position of the JunD peptide used for the crystallization of menin.

The E366D mutation was found in a MEN1 family and localized in the MLL1 and JunD binding pocket (Poncin *et al*., 1999). However, E366D did not affect interactions with MLL1/MLL2 or JunD complexes, which suggests that the functional effects of this glutamate-to-aspartate conversion are rather subtle and missed in our assay system. Similarly, R322H and E408Q isolated from a sporadic glioma and non-small cell lung cancer patient, respectively, did not affect MLL1/MLL2 or JunD complexes, which suggest that these effects are either minimal or passenger rather than driver mutations for these pathologies.

Taken together, the nuclear interactome data of our *MEN1* mutant set indicates a pathological role for six out of nine menin mutations tested. In two recent studies *MEN1* gene missense mutations were analyzed in different *in silico* approach to predict unstable menin mutants (Biancaniello *et al*, 2022; Caswell *et al*, 2019). In contrast, we combined *in silico* predictions for stable menin mutants with transient and stable expression in HeLa cell and quantitative mass spectrometry to identify deficient functions of these mutants. Our functional analyses support the notion that restoration of menin-MLL1 interactions is a potential treatment goal in MEN1 and *MEN1*-related endocrine tumors (Dreijerink *et al*., 2017b).

The menin protein is under strict quality control mechanisms operating in the cytoplasm as indicated by its protein interactors in this cellular compartment. Eight of the top 10 interactors of WT menin either belong to the protein chaperone or the ubiquitination pathways (Figure 4A). It is unlikely that this results from an overexpression artefact as all menin-GFP proteins are expressed to (much) lower levels than endogenous menin (Figure 2A). It is also unlikely that interaction with the protein chaperone or ubiquitin pathways depend on the GFP moiety as similar experiments with the cytoplasmic fraction of other transcription factors do not identify these protein chaperone or the ubiquitination proteins (Antonova *et al*, 2018). Strikingly and as expected, the L168P and L264P instability mutants display increased interactions with components of chaperone and ubiquitination pathways. In contrast the other menin mutants do not display such increased interactions, which indicates that they are similarly stable as WT menin. We note that the E366D menin mutant displayed a strong interaction with the nuclear pore component NUP210 (Figure 4B). NUP210 has recently been proposed to also act as a mechanosensor connecting the cytoplasmic components to the nucleus to control heterochromatin formation (Amin *et al*, 2021). Possibly, loss of tumor suppressor activity by the E366D mutation relates to an acquired function related to NUP210 binding.

Genome wide DNA occupation analyses showed that menin and JunD are present together at many sites of active transcription (Figure 5). While the menin-JunD interaction may not be critical for the molecular pathogenicity of *MEN1* gene mutations, this remarkable co-existence remains intriguing and could still be relevant for menin function, besides endocrine tumorigenesis. It should be noted that cell-type specific effects cannot be ruled out, as due to lack of a suitable endocrine cell line, we used human cervical carcinoma-derived HeLa cells for our experiments. Similar to menin, the JunD transcription factor is broadly expressed across many tissues and we expect that our observations in HeLa cells are valid for many different cell and tissue types. We found that the menin binding pocket mutants R52G, E255K and E359K have differential effects on MLL1/MLL2 and JunD interactions, which translate into differential genomic binding patterns. This indicates subtle differences in MLL1/MLL2 and JunD binding, which are consistent with the structural data on MLL1 and JunD peptide binding (Figure 7). In contrast, the E366D mutation observed in a MEN1 family was predicted to affect MLL1 and JunD binding, but it had little effect on both (Figure 3B), which may be explained by the conservative nature of the E366D mutation.

Taken together, our findings underline the effects of *MEN1* gene mutations in both familial and sporadic tumors of endocrine origin on the interaction of menin with MLL1 and MLL2 in COMPASS-like histone H3K4 methyltransferase complexes and with JunD-containing transcription factors. In addition to menin’s role as a tumor suppressor in MEN1-associated endocrine tissues, the *MEN1* gene has pro-oncogenic activity in the hematopoietic system. A role for AP-1 transcription factors like JunD in MLLr and NPM1 mutant acute leukemias has not been explored yet, but the first clinical trial results with menin-MLL inhibitors combined with our menin mutant results motivate future studies of these inhibitors on the menin-JunD interaction and its possible functional involvement in acute leukemias.

## Acknowledgements

The research of EOG and HThMT is financially supported by the grants from the Deutsche Forschungsgemeinschaft (DFG, German Research Foundation) by 192904750-SFB 992, SFB850 subproject B9, and TI688/1-1. KD received special support from DFG-funded SFB850 for a short sabbatical stay in Freiburg. We thank Christoph Peters for support and Tanja Bhuiyan, Timothy Chan and Saiful Islam for critical reading of the manuscript and all other members of the Timmers lab for discussions and their support.

## Figure legends of Supplementary Figures

Figure S1

Volcano plots of interactors of mutant menin-GFP isolated from nuclear extracts. The mutant menin-GFP proteins is indicated by black dot, whereas subunits of MLL1/MLL2 complexes are indicated in red and JunD-containing complexes in blue.

Figure S2

Volcano plots of interactors of mutant menin-GFP isolated from cytoplasmic extracts. The mutant menin-GFP proteins is indicated by black dot. Proteins involved in ubiquitination are indicated in green and protein chaperones in orange.

